# Generative AAV capsid diversification by latent interpolation

**DOI:** 10.1101/2021.04.16.440236

**Authors:** Sam Sinai, Nina Jain, George M Church, Eric D Kelsic

## Abstract

Adeno-associated virus (AAV) capsids have shown clinical promise as delivery vectors for gene therapy. However, the high prevalence of pre-existing immunity against natural capsids poses a challenge for widespread treatment. The generation of diverse capsids that are potentially more capable of immune evasion is challenging because introducing multiple mutations often breaks capsid assembly. Here we target a representative, immunologically relevant 28-amino-acid segment of the AAV2 capsid and show that a low-complexity Variational Auto-encoder (VAE) can interpolate in sequence space to produce diverse and novel capsids capable of packaging their own genomes. We first train the VAE on a 564-sample Multiple-Sequence Alignment (MSA) of dependo-parvoviruses, and then further augment this dataset by adding 22,704 samples from a deep mutational exploration (DME) on the target region. In both cases the VAE generated viable variants with many mutations, which we validated experimentally. We propose that this simple approach can be used to optimize and diversify other proteins, as well as other capsid traits of interest for gene delivery.

## Introduction

Protein engineering is of increasing importance in modern therapeutics. Natural proteins evolved to satisfy biological requirements for their organisms, however, some of these properties are also of use for therapeutic purposes. The Adeno-associated virus (AAV) makes a great example: this non-pathogenic virus is currently a primary candidate to be used as delivery vector in gene therapy, and is the delivery mechanism of choice in FDA-approved gene therapies (Dunbar et al., 2018; Grimm et al., 2008). However, as a naturally occurring virus, it hasn’t adapted to satisfy all of the properties that are desirable for a vector. For instance, many human sera have neutralising capability against the virus (Calcedo et al., 2009; Mingozzi et al., 2015). In order to alleviate these problems, a first step would be to generate a diverse set of sequences that maintain the ability to assemble and package DNA. We assume that more distant and diverse functional sequences compared to the circulating populations of natural variants increase the probability of finding sequences that for instance, can avoid natural immunity (Bryant et al., 2021). However, finding sequences of high diversity can be challenging for reasons that we outline below.

High-throughput DNA synthesis technologies allow for direct design of thousands of sequences for experimental validation (Adachi et al., 2014; Bryant et al., 2021; Ogden et al., 2019). Previously, high-throughput diversification was largely performed through random mutagenesis and DNA shuffling (Sarkisyan et al., 2016; Yang et al., 2019). These methods have the drawback of extremely low yield as a large majority of random mutations result in non-functional capsids in the selection phase (Bryant et al., 2021; Ogden et al., 2019). On the other hand, low throughput “rational design” approaches were used for precise design of a small number of variants that generally yield a higher proportion of functional variants (Havlik et al., 2020; Maurya et al., 2019). However, direct high-throughput synthesis scales better than what is reasonable for rational design. This provides an opportunity for modern computational approaches to step in as an intermediate where many sequences can be automatically designed in high throughput, with far better yield than random mutations.

Classical computational approaches to protein design based on biophysical first principles have enjoyed substantial success in the past decade (Huang et al., 2016). However, more recently, data-driven machine learning efforts for protein engineering have seen a rapid uptick, targeting protein structure and (to a lesser degree) function prediction (Alley et al., 2019; AlQuraishi, 2019; Hiranuma et al., 2021; Hopf et al., 2017; Marks et al., 2011; Norn et al., 2021; Rao et al., 2019; Senior et al., 2020). These efforts often make use of large protein databases to learn general rules that govern protein sequences within families and are at times followed up by finetuning on specific sequences. For function prediction, available mutational scanning datasets can then be used to evaluate the ability of such models to predict the effects of small perturbations(Hopf et al., 2019; Riesselman et al., 2018).

In the case of AAV, the standard approaches both in computational design as well as transfer learning through ML can come with drawbacks. The AAV capsid consists of a 60-subunit assembly of viral protein (VP) monomers, making accurate biophysical modeling challenging. Previous attempts in rational computational design of AAVs include building a phylogenetic tree and using that as a reference to reconstruct ancestral viruses(Zinn et al., 2015), though the resulting sequences did not display improved immune evasion. On the other hand, computational evolutionary (unsupervised) methods for predicting function are unknown to be effective in this setting as the number of known homologues in the dependo-parvovirus family is relatively small (<1e3).

We recently conducted a high-throughput study with a supervised ML approach to diversify AAV capsids (Bryant et al., 2021). This method shows promise for supervised design of AAV capsids It builds upon mutational scans in the sequence space near the original capsid. However, as a supervised method, it makes no use of evolutionary information. Similarly, Marques et al. use supervised methods to predict AAV capsid assembly from plasmid library data (Marques et al., 2021). In this work, we investigate the power of using simple unsupervised methods, including Variational Auto-Encoders (VAE) to extract useful information out of evolutionary data together with deep mutational explorations. In this case, a deep mutational exploration consists of randomly and additively sampled variants with up to 23 substitutions. This is a subset of the training set used in (Bryant et al., 2021) restricted to substitutions only, and with training samples that we estimate as likely to assemble and package (no negative labels). In particular, we are interested in diversifying the AAV capsid using this information. The promise of combining evolutionary and assay-labeled data has also been observed in other contexts (Hsu et al., 2021; Wittmann et al., 2021).

Previous work has shown that VAEs are effective in capturing the effects of mutations when trained on evolutionary data, but most of these results are applied to small edit-distances (Riesselman et al., 2018). They have also been shown to learn relevant latent space structure that captures phylogenetic relationships and propose new proteins (Ding et al., 2019; Greener et al., 2018). In this work, we focus our efforts on generating variants of AAV2 VP3 protein that are able to successfully assemble and package (we refer to this trait as viability), a prerequisite for any downstream engineering task, using evolutionary and experimental data. We select a 28 amino acid window near the heparin binding site, for which we have collected data previously and which is known to be of immunological significance (Tseng and Agbandje-McKenna, 2014). The MSA consists of 564 sequences from AAV2-related strains. The experimental data we use consists of 36,562 variants of AAV2 (substitution only) mutants of which 22,704 we classified as viable and were designed by an in-silico random model or based on a single site mutation model (see supplement for details).

We investigate three partitions of data, and two unsupervised models. Our model consists of an independent-sites model (IS) as baseline, and a small VAE model (1e5 parameters) with two latent variables with the same architecture as in (Sinai et al., 2017). Our datasets are (i) Evolutionary data (MSA) alone (n=564) (ii) Evolutionary data supplemented 667 randomly generated viable variants (MSAr) from a deep mutational exploration (n=1,231) and finally (iii) Evolutionary data augmented by 22,704 viable (MSA+) mutants 1-21 mutations from wildtype generated at random or an additive model (n=23,268). Producing variants without access to any labeled data would have been risky as the MSA consisted of only a few hundred sequences. Having the deep mutational exploration data allowed us to also test the model’s score (calculated by reconstruction probability) on a set of 7150 holdout variants (both viable and non-viable), to ensure that the VAE captures information about the variants. Both models are trained in an unsupervised manner.

To generate variants with the IS model, we constructed a position weight matrix (PWM) for the sequence and sampled each column proportional to the normalized probability of each amino acid occurring (ignoring gaps). We then sorted the variants according to their log reconstruction score and selected the top 1250 variants for each model-dataset pair (7500 in total for both models). To generate sequences from the VAE, we induced posterior distributions by generating a grid of radial latent coordinates for the VAE centered at the center of mass of the evolutionary data, and then decode the variants (i.e., emitting conditional distributions using generators seeded with the latent code). We then sampled this distribution similarly to how we sampled the IS PWMs, and we always included the maximum likelihood sequence for every conditional (local) PWM. This approach was used to investigate the ability of the VAE to interpolate between known viable regions in the latent embedding. We sorted the variants based on the model score and included the top 1250 variants per pair (See Fig 1 for a schematic).

**Figure 1.**
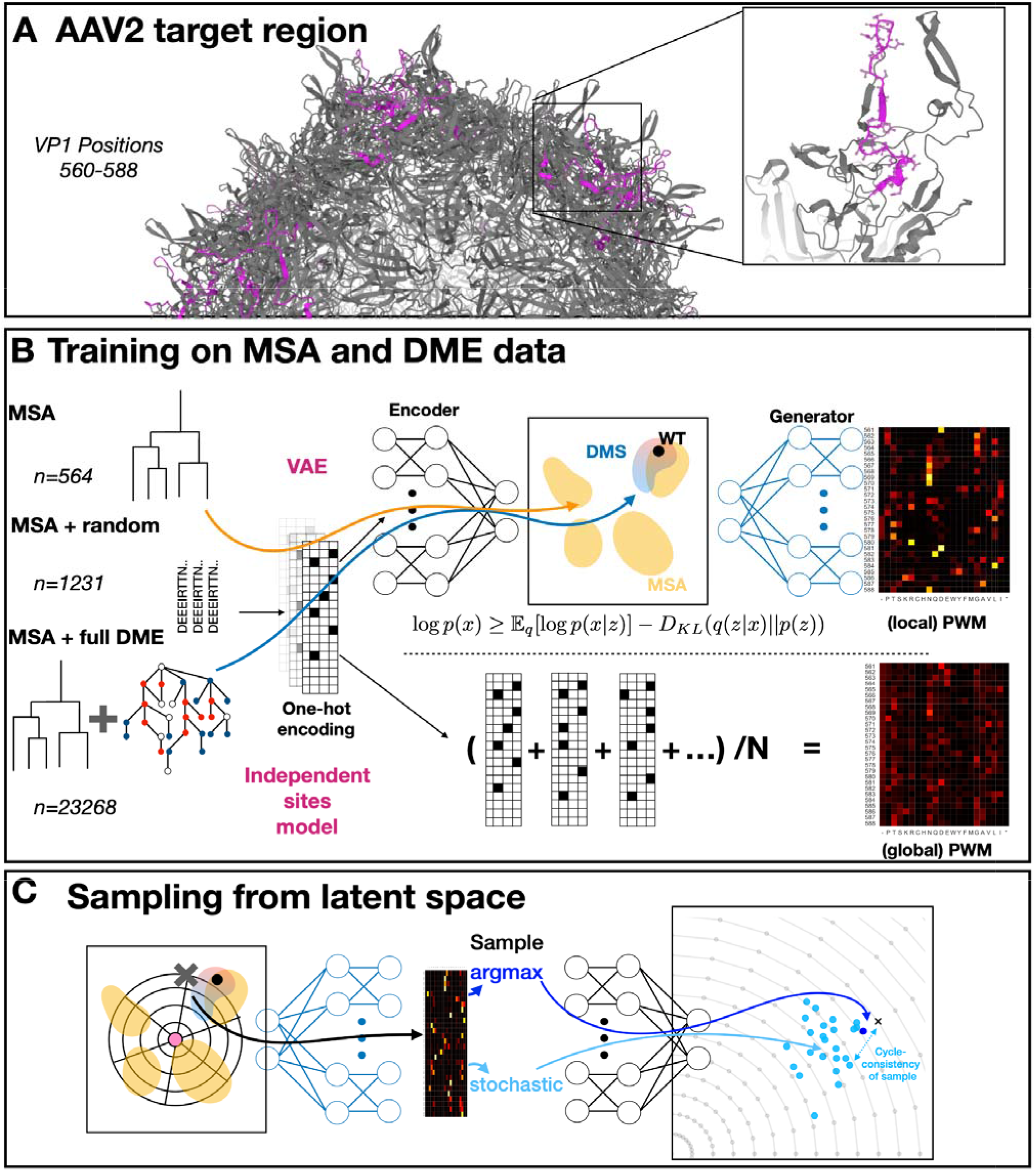
Overview of the generative process. (A) Targeted region (in purple) between VP1 positions 560-588 includes buried and exposed capsid segments with a variety of secondary structures and is positioned at a 3-fold axis contact point crucial for cell tropism, immune evasion and assembly. (B) Our training process on MSA+DME data. The data is one-hot encoded and passed through a VAE as well as the IS model. Both methods generate PWMs from which samples are drawn stochastically with a fixed temperature. VAEs are optimized against an objective that jointly minimizes KL-divergence between encoding q(x|z) and a latent standard normal prior p(z) with the expected log reconstruction error of the output. This is known as the Evidence Lower BOund (ELBO), see (Kingma and Welling, 2013)(C) Sampling from the VAE latent space involves starting at latent coordinates, inducing a conditional PWM, and then sampling from it stochastically, as well as by taking the argmax per position. As a sanity check we passed each sample through the encoder and confirmed that they are generally correlated to the original point.

We compared the experimental production score of our training set variants with log reconstruction probability of the model in generating those variants. On the holdout samples, we found high correlation between IS and VAE model scores (Spearman *ρ*~0.95 for all models trained on the same training set), that is the models highly agree. We also found moderate correlation for the model scores and experimental holdout data (Spearman *ρ* (i) IS: 0.41, VAE:0.47, (ii) IS: 0.48, VAE:0.5, (iii) IS, VAE = 0.55). We deemed this sufficient to use the models for in-vitro validation. Our cloning and production assays were done exactly as described in (Bryant et al., 2021) with additional details explained in the SI. In short, we calculate a selection score *s_i_* = *log*_2_(*f_i_^virus^* / *f_i_^plasmid^*) where *f_i_* denotes the frequency of a variant in plasmid or virus pool measured through next generation sequencing (NGS). Overall, our experiments show excellent correlation among replicates (*r*^2^ = 0.98, see SI Fig 1).

## Results

Our primary objective in this study was to explore the sequence space for diverse and viable variants in response to different models and datasets. We classify our viral production into three categories: Viral variants likely arising from dysfunctional genes (observed in production due to low frequency cross-packaging into viable capsids) termed *non-viable*, viral variants likely capable of generating functional assemblies termed *viable*, and viral variants that are present at very high counts, termed *viable*+ (See more detailed discussion of this classification in the SI).

Encouragingly, we found more than 100 viable variants with 25 or more substitutions in the 28-aa region, a majority of which came from the VAE designs. Of the top 10 variants produced, 9 were proposed by VAEs, and 1 by IS-MSA+. Of the top 100 scoring variants produced, 70 were proposed by the three VAE models, and the rest were proposed by IS-MSA+ (see Supp Fig 2). Overall, the IS model trained on the MSA+ dataset performed best in terms of yield with 92.6% of proposed sequences being viable (See supplementary table 1) however the yield was a result of picking the top (per-site) mutations present in the training data, and the farthest designed variant in this set was only 5 mutations away from the WT. The IS model’s performance deteriorated with the removal of the large mutational scan data. For the VAEs the yield was somewhat consistent among the datasets, with the VAE-MSAr pair performing marginally better than the other methods. Even the shallow MSA dataset alone was sufficient to produce a large number of viable variants with many mutations away from the reference sequence (Fig 2E). This is in part an indication of modular interchangeability among dependo-viruses, despite significant sequence-level variation (See SI Fig 2), but as indicated above, IS models were insufficient in generating these viable variants.

**Figure 2.**
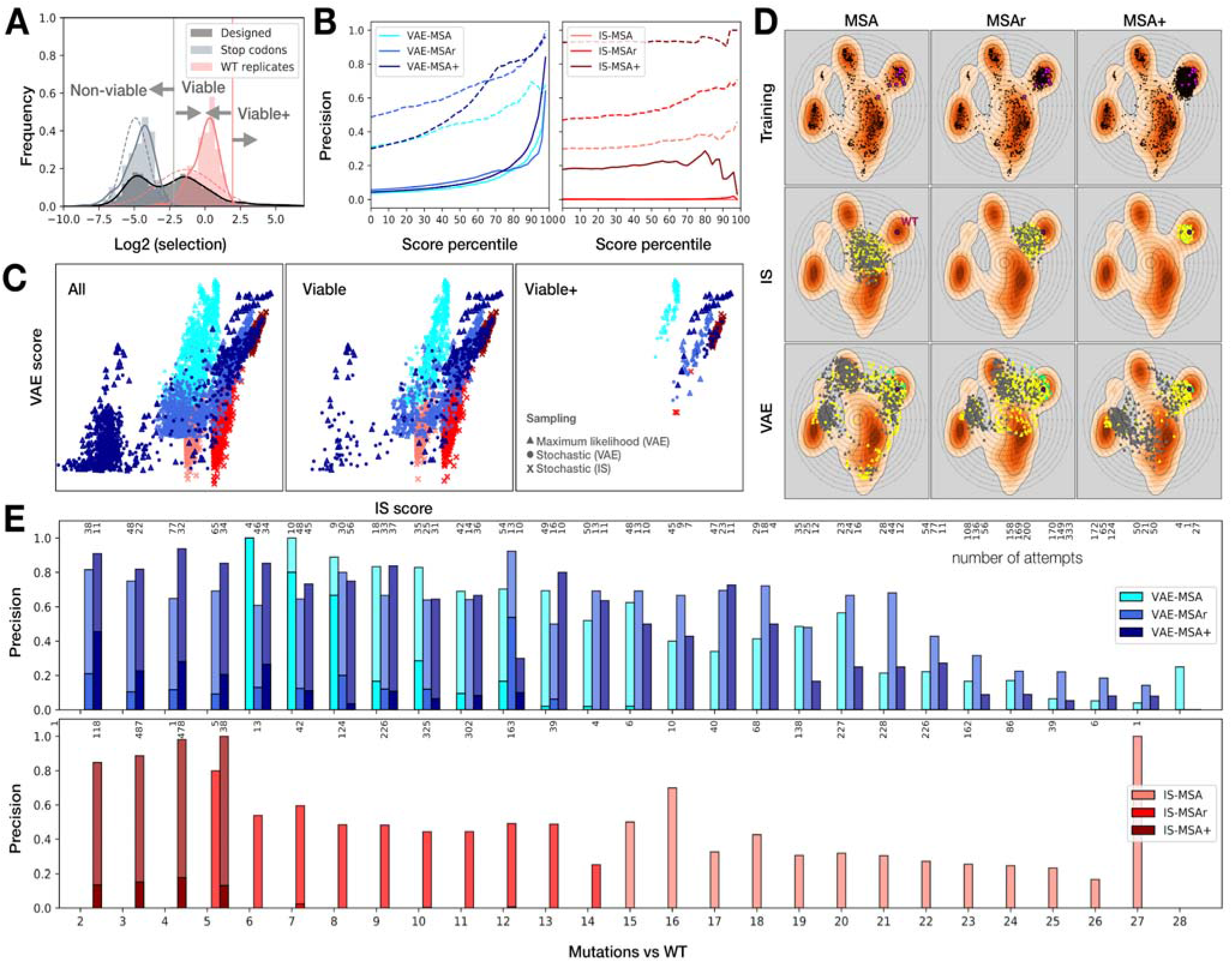
Successful generation of variants with VAE and IS models. (A) Classification of variants into non-viable, viable and viable+ is done by conservative decision bounds computed based on the distribution of the two Gaussian distributions (approximated by dashed lines), representing viable and non-viable distributions (see supplement for details). These are largely in agreement with WT and stopcodon replicates from the Bryant et al. study shown for comparison. Variants with selection score above 3 standard deviation of WT average and 97 percentiles of the Gaussian fit for successful producers are classified as viable+. (B) The relationship between model-denoted score and probability of success. The x-axis denotes the percentile score for the variant based on the model’s evaluation. Dashed lines denote viable fraction, solid lines denote viable+. Precision denotes fraction successful by the given criteria (viable or viable+). (C) Comparison between the IS and VAE scores for variants. The first panel includes all variants, the second panel viable variants, and the third panel viable+. Triangles denote argmax sampling, circles denote stochastic sampling from VAE posterior, x denote stochastic sampling from IS prior. Colored as in part B. (D) Distribution of sampled variants projected onto the VAE-MSA latent dimensions. Orange contours are density plots for evolutionary data alone (points in the first panel). WT denoted by a purple dot. Magenta points show projected location of AAV1-12 onto the same space (top row). Grey(non-viable), yellow(viable), Green (viable+) points in the bottom two rows show the projection of experimental samples onto the latent coordinates. Circles are stochastic samples, whereas triangles are maximum likelihood (argmax). (E) Distribution of samples with viable (lighter bars) and viable+ (darker bars) variants at each distance sampled by each algorithm. The number of attempted samples is denoted at the top of each column. Precision denotes the fraction of those attempts that resulted in success.

A key observation however is that both the IS and VAE models performed significantly better in terms of yield when “fine-tuned” through augmentation with the mutational scan data. As shown in Fig 2B, adding more data increases the quality of the IS model substantially. For the VAE however, it appears that the intermediate MSAr dataset is comparable to the full MSA+ for model quality despite having 10-fold fewer samples in its training data. Comparing how the two models score variants we find that every viable+ variant scores highly with both models (although a large majority are sampled by a VAE), however, some viable variants proposed by the VAE score poorly with the IS model (when trained on the same dataset, Fig 2C). Curiously, high-scoring VAE variants were far more likely to be in the viable+ category (Fig 2B, C).

Overall, our generative models manage to produce a significant number of viable capsids with more than 50% of positions substituted, with multiple viable variants that had substitutions at almost every position. This is not a trivial task: Most random changes in the amino-acid space within this region result in a non-functional capsid (Bryant et al., 2021).

## Discussion

We mentioned in previous segments that the predictive accuracy of the IS and VAE models is very similar. But the sampling schemes introduced here exhibit drastic differences in both the quality and the diversity of samples between VAEs and the IS model. Yet, both methods perform decently as generative methods, with far better yield than random sampling as seen in (Bryant et al., 2021; Ogden et al., 2019). The performance of the IS model is consistent with previous studies where independent effect terms explain a large fraction of trait variability (Otwinowski et al., 2018). Furthermore, our VAE model can be thought of as a mixture model, with each set of initial coordinates emitting a single sequence profile, within which additive effects explain the majority of variation (Dauparas et al., 2019; Marshall et al.).

While IS models and VAEs share similarities, VAEs seem much better suited to use for generative interpolation. We are encouraged by the observation that even a shallow MSA is sufficient to generate interesting and high-performing variants with the VAE, and furthermore, adding only a few hundred randomly sampled variants (achievable through simple error-prone PCR instead of direct synthesis) drastically increases the performance of these models. This suggests an experimental and computationally accessible framework can be used to generate thousands of viable variants, both for AAVs and other proteins of interest.

This work stands to benefit from further investigations in multiple directions. First, in training these models, we ignored the full sequence (i.e., background) in which these AA substitutions were being made. We chose a segment that could be directly synthesized and was of sufficient challenge and clinical relevance. However, at least the VAEs have a clear possibility of learning from the context and improving their predictions as a result. Furthermore, for both IS and VAE models, we use a fixed temperature for sampling variants. It would be interesting to tune the temperature as a method of generating further diversity. The IS model can also be augmented with clustering to generate local sequence profiles, increasing diversity in the samples. In addition, while we kept the latent dimensions of the VAE limited to 2 to simplify interpolation, it is a tunable hyperparameter, and we expect that higher capacity models could be used to perform more sophisticated sampling with better-fit models. Our study is a proof-of-concept for augmenting MSAs with experimental measurements with generative models, and was conducted in a fully unsupervised manner. However, it is also possible to conduct training in a semi-supervised fashion, where available labels are used to help the VAE learn relevant latent representation. Finally, generative models like those in this study can be paired with supervised methods to optimize sequences toward particular traits (Sinai and Kelsic, 2020; Sinai et al., 2020). Brookes et al. develop such a method which uses a VAE together with supervised oracles to optimize GFPs *in-silico* (Brookes and Listgarten, 2018; Brookes et al., 2019). Our study indicates an increased probability of success for these methods to be applicable in AAV design. We show that even without the need for deep mutational data, VAEs have potential as generative approaches for designing proteins that are hard to model structurally (like capsids). This should open avenues for low-cost and fast computational design with much higher yield than random mutagenesis, in a manner accessible to many wet lab scientists.

## Acknowledgements

We thank Javen Tan with assistance in software development. We also thank Debora Marks for assistance with the alignments. We thank David Ding and Martin Nowak for related discussions. We thank Jakub Otwinowski, Elina Locane, Farhan Damani, Michael Stiffler, Jeffrey Gerold from Dyno Therapeutics for helpful comments on the manuscript. We thank members of the Church lab for their support, in particular Surge Biswas, Gleb Kuznetsov, and Pierce Ogden. This work was supported by the Wyss Institute.

## Code and Data Availability

Code and data for this paper can be accessed here: https://github.com/churchlab/Generative_AAV_design

## Supplementary material

**Supplementary Figure 1.**
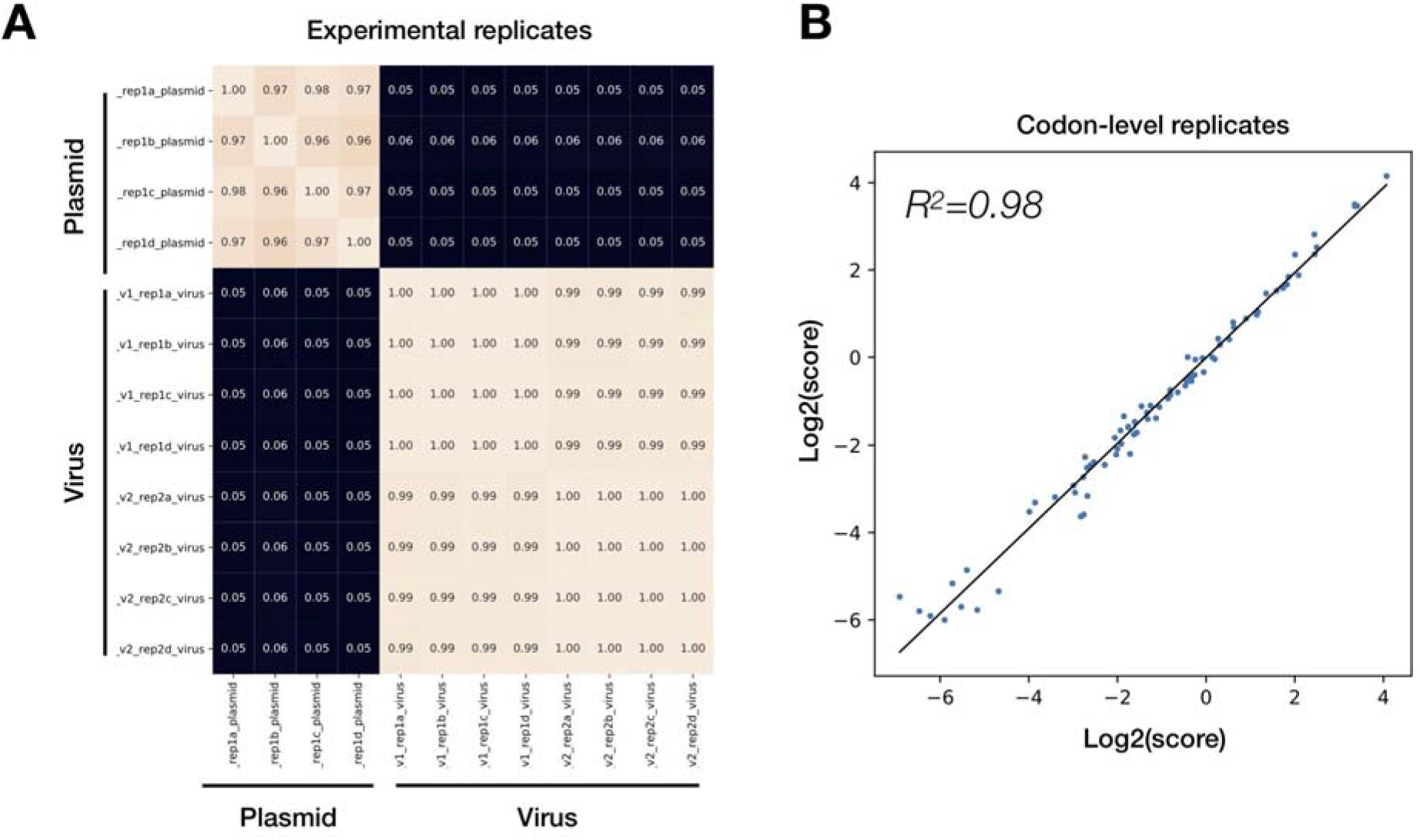
Experimental reproducibility. (A) We show correlation between counts among 4 separate plasmid and 4 separate virus production replicates (each with two PCR replicates). As expected, plasmid and viral counts per variant show no correlation, whereas there is excellent correlation within virus and plasmid replicates. (B) We included n=196 codon-level controls (different codons but same amino-acid sequence) corresponding to n=78 unique amino-acid sequences in our synthesis, confirming that there is excellent correlation between codon-level replicates.

**Supplementary Figure 2.**
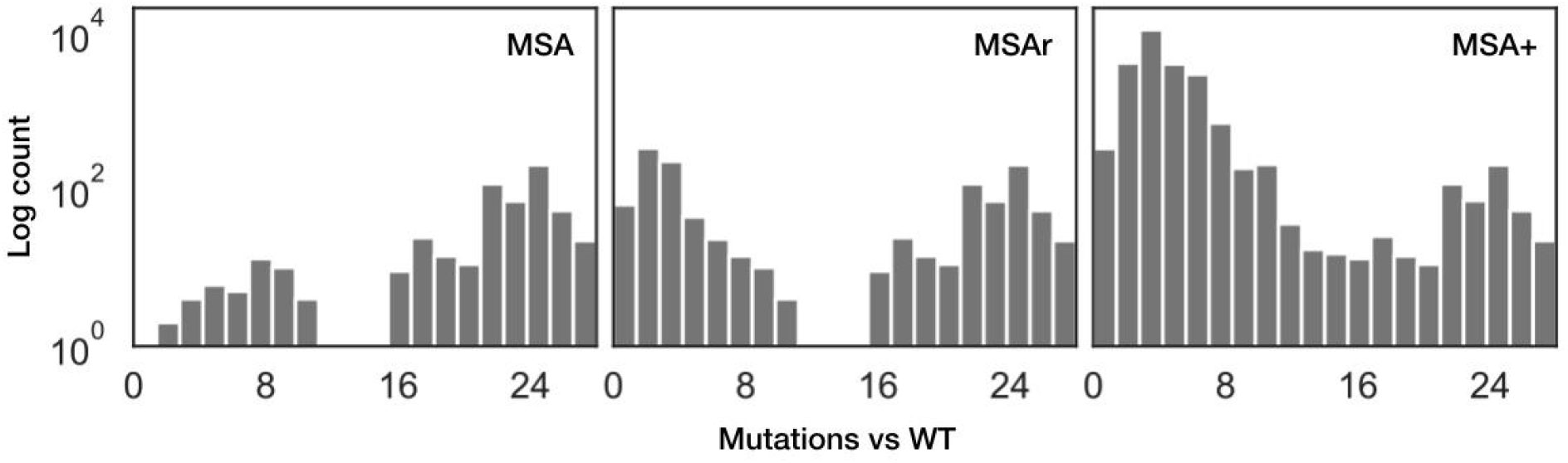
Distribution of substitution counts in training data. A majority of MSA variants are more than 16 mutations away from AAV2. On the other hand, a majority of the MSA+ falls within 5 substitutions of the reference variant.

**Supplementary Figure 3.**
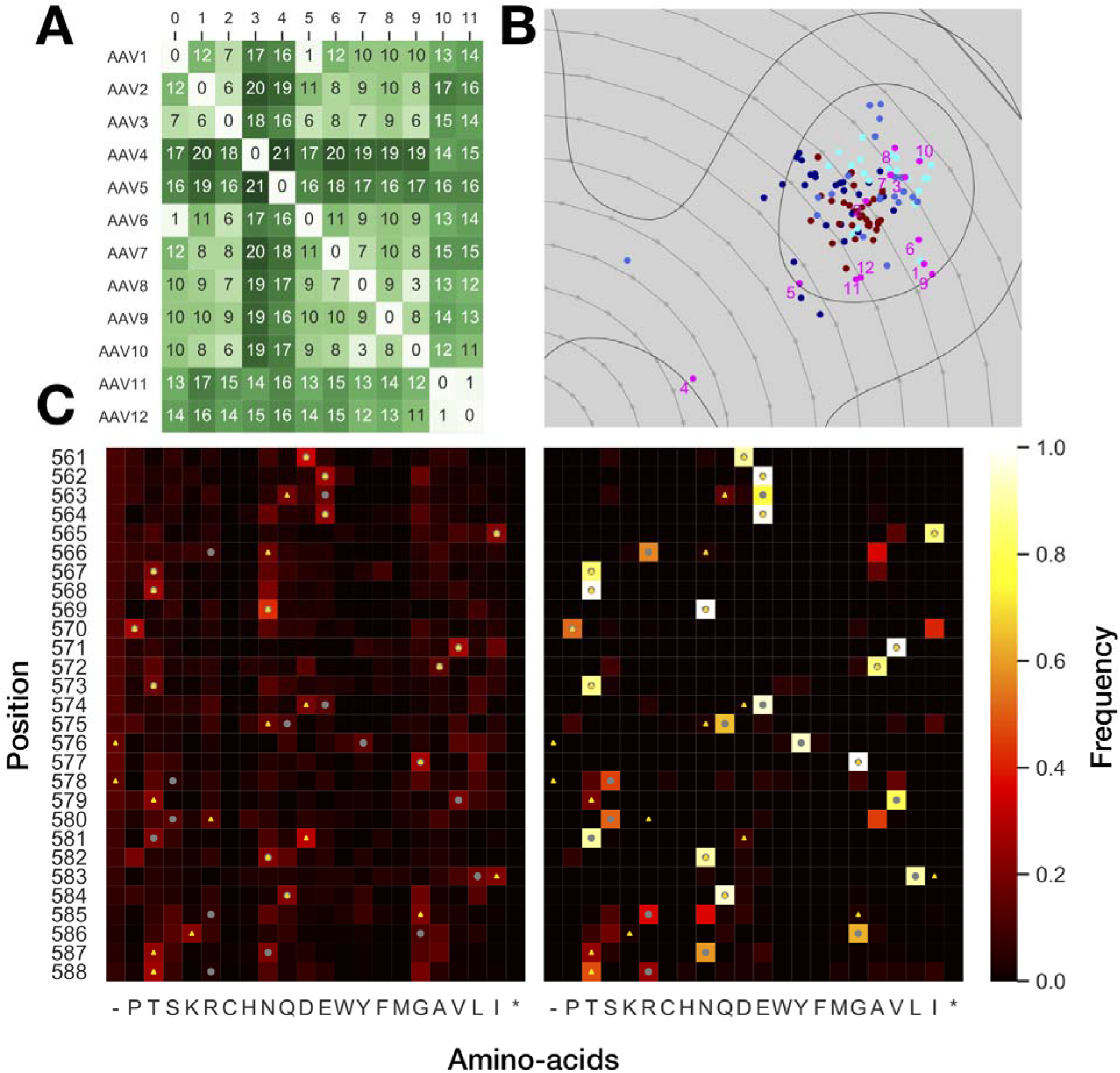
(A) Levenshtein edit-distance of the 560-588 segment among common AAV serotypes. (B) Projection of the top 100 produced variants onto the VAE latent space, colored as in Figure 2 for each method, along with the AAV serotypes for reference. (C) Left PWM for the IS model, with wildtype positions marked with a grey dot, and the maximum row-wise value marked by a yellow triangle. Right: PWM for top 100 variants. Star (*) represents the stop codon, and (–) indicate gaps in the alignment.

**Supplementary Table 1.** Number and proportion of variants designed by each method resulting in viable and viable+ samples.

|  | MSA | MSAr | MSA+ |
| --- | --- | --- | --- |
| **IS** | Viable+: 0 (0%)<br>Viable: 369 (29.8%)<br>Total: 1237 | Viable+: 3 (0.2%)<br>Viable: 583 (46.9%)<br>Total: 1244 | Viable+: 116 (10%)<br>Viable: 1040 (92.6%)<br>Total: 1122 |
| **VAE** | Viable+: 34 (2.7%)<br>Viable: 385 (31.0%)<br>Total: 1242 | Viable+: 44 (3.5%)<br>Viable: 587 (47.3%)<br>Total: 1242 | Viable+: 42 (3.4%)<br>Viable: 370 (29.8%)<br>Total: 1243 |

## Supplementary information

### Library Production

Libraries were produced precisely as and in parallel with the experiments described before in (Bryant et al., 2021) (see methods). We also used the same original additive training set as our starting point to construct our training library, described therein. We successfully cloned 7446 out of the 7500 (99.28% yield) of our designed sequences as plasmids but only analysed the 7409 for which we sampled at least 100 plasmids in the cloned pool.

### Viral production classification

We have observed previously that production assays tend to follow a bimodal mixture of Gaussian distributions ((Bryant et al., 2021), see supplementary Fig 1), with scores in the first mode closely resembling those of the negative controls.

We numerically fit the distribution of scores within our library with a Gaussian mixture-model (GMM), which yields *μ_gmm_^pos^* = −1.35, *μ_gmm_^neg^* = −4.95 and *σ_gmm_^pos^* = 2.04, *σ_gmm_^neg^* = 0.85.

Furthermore, we compare our fit results against wildtype (WT) produced (which is expected to score higher than the typical variant as many mutations are deleterious) and stop-codon controls produced in a separate experiment. There we had *μ_wt_* =-0.80, *μ_stop_* =-5.47 and *σ_wt_* = 0.91, *σ_stop_* = 1.03. Using these two sets of approximations for scores we bin the data into three categories: Non-viable variants that do not package DNA efficiently (similar to stop-codon in performance), viable variants (between non-viable and highest WT replicate score), and viable+ which we categorize as extremely good capsid producers.

We calculate the boundary between viable and non-viable as follows. We approximate the distribution of WT and stop-codon controls by Gaussian distributions and calculate *b_viable_* = (*μ_wt_* − 2*σ_wt_* + *μ_stop_* + 2*σ_stop_*)/2. For the better than WT boundary, we calculate *b_viable+_* = (*μ_wt_* + 3*σ_wt_*). These boundaries yield the following values for the cumulative distribution function of our GMM fits: *CDF* (*b_viable_*; *GMM_neg_*) =0.999, *CDF* (*b_viable_*; *GMM_pos_*) = 0.335, *CDF* (*b_viable+_*; *GMM_pos_*) = 0.977. Given this evidence, we believe these are strongly conservative but trustworthy decision boundaries minimizing false positives.

### Sampling from PWMs

We sample variants from PWMs in both IS and VAE model by normalizing frequencies 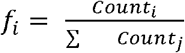in each position and sampling an amino acid at each position (possibly the WT variant) with probability 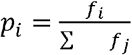.

For Maximum likelihood samples, we take the most likely amino acid for each position. We don’t use a temperature parameter (that is, we assume a fixed temperature).

### Alignments

We performed a Jackhmmer search (2015 update (Finn et al., 2015)) against UniprotKB for the VP positions 200-735 yielded a clear clustering of high-scoring hits based and were pruned based on qualitative inspection of the significance scores. We then extracted the 28 amino acid regions of interest from this data, resulting in 564 sequences.

